# Power and limits of selection genome scans on temporal data from a selfing population

**DOI:** 10.1101/2020.05.06.080895

**Authors:** Miguel Navascués, Arnaud Becheler, Laurène Gay, Joëlle Ronfort, Karine Loridon, Renaud Vitalis

## Abstract

Tracking genetic changes of populations through time allows a more direct study of the evolutionary processes acting on the population than a single contemporary sample. Several statistical methods have been developed to characterize the demography and selection from temporal population genetic data. However, these methods are usually developed under the assumption of outcrossing reproduction and might not be applicable when there is substantial selfing in the population. Here, we focus on a method to detect loci under selection based on a genome scan of temporal differentiation, adapting it to the particularities of selfing populations. Selfing reduces the effective recombination rate and can extend hitch-hiking effects to the whole genome, erasing any local signal of selection on a genome scan. Therefore, selfing is expected to reduce the power of the test. By means of simulations, we evaluate the performance of the method under scenarios of adaptation from new mutations or standing variation at different rates of selfing. We find that the detection of loci under selection in predominantly selfing populations remains challenging even with the adapted method. Still, selective sweeps from standing variation on predominantly selfing populations can leave some signal of selection around the selected site thanks to historical recombination before the sweep. Under this scenario, ancestral advantageous alleles at low frequency leave the strongest local signal, while new advantageous mutations leave no local footprint of the sweep.

## Introduction

Several evolutionary processes (such as migration, selection or drift) can change the genetic make-up of populations through time. Thus, patterns of genetic diversity can inform us about the evolutionary history of the populations (see Pool *et al.* 2010, for a review). However, observing the genetic diversity changes through time (instead of at a single time point) can provide more precise information about the evolutionary processes in action.

Since the beginning of the 20th century, researchers have used repeated observations of hereditary characters in the same populations (e.g. color patterns in *Diabrotica soror*, Kellogg and Bell 1904) or subfossil records (e.g. banding patterns in *Cepaea* snails, Diver 1929) to study evolution. An emblematic example is the time-series data on the frequency of the *medionigra* phenotype in a population of the moth *Callimorpha dominula*, which inspired the discussion on the prevalence of selection over drift between Fisher and Wright (Fisher and Ford 1947; Wright 1948) and has continued to offer insight into the evolutionary process through recent re-analyses (e.g. Foll *et al.* 2014, and references therein).

The technological advances in molecular genetics have allowed these temporal studies to switch from Mendelian characters to polytene chromosomes (e.g. Dobzhansky 1943), isozymes (e.g. Yamazaki 1971) and, eventually, to high throughput DNA sequencing (e.g. Frachon *et al.* 2017). Indeed, molecular genetics has opened the door to the study of short-generation-time microorganisms (Biek *et al.* 2015), ancient samples (e.g. subfossil samples, museum and herbaria specimens; Leonardi *et al.* 2017) and experimental populations (Schlötterer *et al.* 2015), allowing for an increasing availability of temporal population genetic data.

Temporal population genetic data allow to study the change of allele frequencies through time. In the absence of migration, mutation and selection, these changes are the product of genetic drift. As such, they can be used to estimate the effective population size, *N*_e_, either with moment based (e.g. Krimbas and Tsakas 1971; Nei and Tajima 1981; Waples 1989) or likelihood-based approaches (e.g. Anderson *et al.* 2000; Williamson and Slatkin 1999). If a sample from the source of migration is available, it is also possible to co-estimate migration and drift with an extension of the likelihood method (Wang and Whitlock 2003). These methods all assume short timescales and low mutation rates, so that no new mutations arrive during the studied period. However, for temporal data over larger scales in which mutations can no longer be neglected, it is also possible to co-estimate substitution rate with effective population size (Drummond *et al.* 2002; Rambaut 2000). Finally, like for migration or mutation rates, it is also possible to make inferences about selection. Two strategies can be followed. First, three or more temporal samples may be used to separate the random component (drift) from the systematic component (selection) in the changes of allele frequencies (e.g. Bollback *et al.* 2008; Buffalo and Coop 2019; Feder *et al.* 2014). Alternatively, if only two temporal samples are available, loci under selection may be detected by an outlier approach (e.g. Goldringer and Bataillon 2004, described in more detail below).

Most of the statistical methods in population genetics, including those mentioned above, have been developed for outcrossing populations. Methods specifically adapted to selfing populations are scarce (Hartfield *et al.* 2017). Nevertheless, many plants reproduce through selfing or partial selfing (see Whitehead *et al.* 2018, for a recent overview), including an important proportion of the species considered in temporal monitoring programs (e.g. appendix A1). Selfing, by increasing homozygosity and linkage disequilibrium, shapes the genetic diversity of populations in a particular way (Golding and Strobeck 1980; Vitalis and Couvet 2001). Notably, it generates repeated multi-locus genotypes that can persist over several generations due to the lack of effective recombination (Jullien *et al.* 2019). The dynamics of adaptation is also impacted by selfing, with a general reduction in the efficacy of selection (e.g. Burgarella *et al.* 2015). In the light of these drastic effects, there is a need for development or adaptation of methods to take into account different mating systems.

Even if we can adapt methods to release the assumption of random mating, distinguishing selection from demography in highly selfing species may be problematic because of the reduced effective recombination due to self-fertilization. Selective sweeps in these populations could involve a genome-wide hitch-hiking effect, which prevents any difference in genetic diversity between the neutral and adaptive regions. If that were the case, even methods adapted to selfing would fail to detect regions under selection, which questions the relevance of temporal genome scans in predominantly selfing populations. On the other hand, a temporal genome scan on a highly selfing *Arabidopsis thaliana* population (selfing rate, *σ* ≈ 0.94) revealed several outlier regions with compelling evidence for the action of selection on them (Frachon *et al.* 2017). Therefore, in the planning of future research, there is a need to understand under which circumstances selfing imposes a limit for the detection of loci under selection.

In this work, we introduce several modifications to the temporal genome scan approach proposed by Goldringer and Bataillon (2004) to take into account partial self-fertilization. Then, by means of simulated data, we evaluate the performance of this method for the estimation of the effective population size (the first step of this genome scan) and the detection of regions under selection under different scenarios of adaptation (from new mutations or from standing variation) and selfing. Our results highlight the importance of taking into account the mating system in the analysis of population genetic data. They also highlight a threshold beyond which loci under selection cannot be detected for highly selfing populations. We applied the approach to a population of the predominantly selfing species *Medicago truncatula* and re-discuss some of the results from Frachon *et al.* (2017) temporal genome scan on *Arabidopsis thaliana*.

## Materials and Methods

### Overview of the genome scan method

In a single isolated population, allele frequencies change through time under the action of selection, which acts upon specific loci, and genetic drift, which acts upon the whole genome. In order to identify loci subjected to selection, we use a procedure inspired from the test for homogeneity of differentiation across loci by Goldringer and Bataillon (2004). The principle is that, in the case of complete neutrality across loci, all sampled markers should provide estimates of genetic differentiation drawn from the same distribution. Assuming a single isolated population, this distribution depends on the strength of genetic drift, that is, on the length of the period, *τ* in number of generations, and on the effective population size, *N*_e_ (see Table A1 for a summary of notation). On the other hand, if some of the studied polymorphisms are under selection or linked to selected variants, we expect some heterogeneity in the distribution of differentiation values, because directional selection induces larger values than expected under the neutral case. The approach we describe uses the expected distribution of temporal differentiation to identify those polymorphisms showing outlier values compared to a neutral expectation. In Frachon *et al.* (2017), we already made some modifications (to account for the uncertainty of the initial allele frequency) to the approach proposed by Goldringer and Bataillon (2004) for the standard (random-mating or haploid) case. Here, we present further modifications for a more general case including partial self-fertilization.

### Estimation of effective population size

The estimated magnitude of drift between two time samples is used as a null model in this temporal genome scan method. Temporal differentiation can be measured by estimating the *F*_ST_ with the analysis of variance approach proposed by Weir and Cockerham (1984). Weir and Cockerham’s (1984) analysis of variance partitions variance within individuals, among individuals within population and between populations, allowing to account for the correlation of allele identity within individuals due to selfing in the *F*_ST_ estimate. Temporal *F*_ST_ can be used to estimate *N*_e_ as 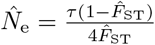 (Frachon *et al.* 2017; Skoglund *et al.* 2014).

### Building null distribution of drift

In order to test the homogeneity between the focal-locus *l* and genome-wide differentiation, the null distribution for single locus 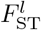 is built through simulations of drift. Each of these simulations consists of the following steps: 1) draw initial allele frequency *π*_0_ of the locus (conditional on data), 2) simulate allele frequency change for *τ* generations (based on 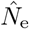), 3) simulate samples by sampling genotypes (genotype frequencies based on 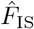) and 4) calculate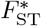 for the simulated sample. The proportion of 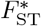 equal or larger than the observed 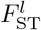 provides an estimate of the *p*-value for the test. A detailed description of these steps follows.

Goldringer and Bataillon (2004) considered the observed allele frequency in the sample as the initial allele frequency in the population, *π*_0_ (from this point subscript 0 indicates values at time *t* = 0). This approach ignores the uncertainty due to sampling. Instead, in Frachon *et al.* (2017), we improved this step by assuming that allele counts observed in a sample of *n*_0_ diploid individuals come from a binomial distribution B(2*n*_0_, *π*_0_), where *π*_0_ is the (unknown) allele frequency in the population. Using Bayes inversion formula and assuming a uniform prior for the allele frequency, this allows to sample from the posterior probability distribution with Beta(*k*_0_ +1, 2*n*_0_ −*k*_0_ +1), where *k*_0_ is the observed count of the reference allele in the sample. However, this assumes that allele copies within an individual are independent samples from the population. In (partially) selfing populations, gene copies are not independent samples, but individuals are. Genotype counts observed in the simulated sample of *n*_0_ individuals can be modelled as coming from a multinomial distribution Mult(*n*_0_, ***γ***_**0**_), where *γ*_**0**_ are the genotype frequencies in the population. Similarly to Frachon *et al.* (2017), assuming the same prior probability for the three genotype frequencies, we sample genotype frequencies in the population from the posterior probability distribution with Dir(*K*_**0**_ + **1**), where *K*_0_ is the observed genotype counts in the sample of the focal locus at time *t* = 0.

Allele frequencies *π_t_* at subsequent generations (*t ∈* [1, *τ*]) were simulated following a binomial distribution as 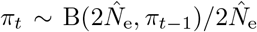, where 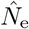 is the genome wide estimate of the effective population size and *π*_0_ is determined by *γ*_**0**_. Simulated genotype counts in sample at time 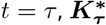, were taken from a multinomial distribution, 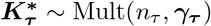, where *n_τ_* is the sample size (in number of diploid individuals) at time *t* = *τ* and ****γ_τ_**** :

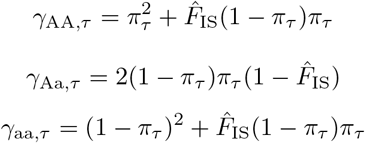

are the genotype frequencies in the populations as a function of the allele frequency *π_τ_* and inbreeding coefficient *F*_IS_ (Haldane 1924), assuming constant selfing rate and using the Weir and Cockerham’s (1984) multilocus inbreeding coefficient estimate from both temporal samples.

Preliminary results revealed that filtering loci according to minor allele frequency (MAF) was required to assure a uniform distribution of *p*-values from neutral sites (Fig. A1). Distribution of *p*-values is important for studies at the genomic scale where thousands of loci are tested (see François *et al.* 2016, for a review). In these studies, a false discovery rate (FDR) is estimated to control for multiple testing and this FDR estimation assumes the uniformity of *p*-values under the null model (Storey 2002). The criterion to filter loci was to have a minimum global MAF: loci with 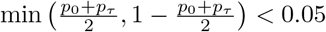 were removed from the dataset. Thus, loci to be tested were chosen based on their genetic diversity, a bias that has to be taken into account in the test. This was done by discarding drift simulations that produced 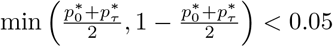, where 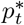 is the frequency of the reference allele in the sample at time *t* in the simulation.

### Simulations

We produced simulated data using the individual-based forward population genetics simulator SLiM 1.8 (Messer 2013). We considered a single isolated population sampled twice, at the beginning and at the end of a time interval of *τ* generations. The population size *N* = 500 (number of diploid individuals) and the selfing rate *σ* were constant through time, and the effective population size was 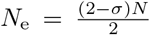 under neutrality (Li 1955; Pollak 1987). The genome was composed by two linkage groups of size 2.5 × 10^8^ base pairs. The neutral mutation rate per base pair was *μ* = 10^−8^ and the recombination rate between base pairs was *r* = 10^−8^. The recombination rate between the two linkage groups was 0.5.

In order to start our simulations with a population at mutation-drift equilibrium, SLiM was run for 20*N*_e_ generations. After that period of neutral evolution, two different selective scenarios were simulated: adaptation from new mutations or from standing variation. In the first case a new advantageous dominant mutation was introduced at *t* = 0 with a selection coefficient *s* at a random position in the genome. In the case of selection on standing variation, a random neutral polymorphic site was chosen at *t* = 0 and a selection coefficient *s* was assigned randomly to one of the two alleles, which also became dominant. The initial conditions of the simulations of selection on standing variation are, therefore, variable. The initial frequency of the allele under selection is known to affect the signal of the sweep: an advantageous allele starting at high frequency is expected to leave a lower signal than one starting at a lower frequency (Berg and Coop 2015; Innan and Kim 2004). This will likely affect the power of our test. We therefore studied the effect of the initial frequency of the advantageous allele in scenarios of adaptation from standing variation with additional simulations, where the site under selection was randomly chosen among those sites with the required allele frequency. The frequency of an allele is expected to correlate with its age (Kimura and Ohta 1973), and older alleles can accumulate more mutations in their neighborhood and recombine with the background variation, which could favour the presence of a local signal of selection (Fig. A2). Because ancestral alleles are older than derived alleles, we also studied the effect of the nature of the allele in the set of simulations of adaptation from standing variation.

In order to ease comparisons across scenarios, the advantageous allele was also set as dominant. Indeed, in outcrossing populations adapting from new mutation, advantageous dominant alleles are expected to spread faster than recessive ones, an effect known as Haldane’s sieve (Haldane 1927). However, this effect is reduced or absent in predominantly selfing populations (Charlesworth 1992; Ronfort and Glémin 2013) and populations adapting from standing variation (Orr and Betancourt 2001).

Simulated populations were sampled at generations *t* = 0 and *t* = *τ*, with samples sizes *n*_0_ = *n_τ_* = 50 diploid individuals. Data for 10 000 polymorphic loci were taken randomly from all polymorphic sites in the sample, except for the locus under selection that was always included in the data.

Different scenarios were considered by exploring values of selfing rate *σ ∈* [0, 0.5, 0.75, 0.8, 0.85, 0.9, 0.95, 0.99, 1], selection coefficient *s ∈* [0, 0.1, 0.2, 0.3, 0.4, 0.5], duration of period of selection *τ ∈* [5, 10, 25, 50, 100, 200] (in generations), type of selection (neutral, new mutation or standing variation) and, in the case of selection from standing variation, whether the locus was chosen randomly among all loci or among the loci with the required initial frequency (*π*_0_ *∈* [0.1, 0.5, 0.9]) and nature (ancestral or derived) of the allele becoming advantageous. The combinations of parameter values were chosen to highlight the patterns studied in this work by creating strong selective sweeps. For each scenario, 100 simulation replicates were performed. Replicates in which the advantageous allele was lost were discarded and replaced by additional replicates.

### Analysis of simulated data

For each simulation replicate, data were analysed as described in the two previous sections. In addition, the effective population size was also estimated from *F*_C_ (following Waples 1989) for comparison. For a given scenario, the true positive rate was estimated from the test results at loci under selection. The false positive rate was estimated from the test results at the neutral loci on the linkage group that does not include the locus under selection. Neutral loci on the linkage group with the locus under selection were used to characterize the footprint of selection due to hitch-hiking by estimating the positive rate as a function of the distance to the locus under selection. In order to quantify the variance of the selection signal, loci were bootstrapped and the 95 % quantile interval for the proportion of positive tests was calculated. Positive tests were defined with an arbitrary threshold of *p*-value<0.001. In order to control for multiple testing, FDR estimates were quantified with *q*-values for each locus, following Storey (2002), with the *qvalue* R package (Storey *et al.* 2019). Manhattan and QQ plots were generated with *qqman* R package (Turner 2017).

As explained in the previous section, the genetic diversity around the selected locus could influence the outlier test in a scenario of adaptation from standing variation in a predominantly sefing species (Fig. A2). Therefore, we examined the structure of genetic diversity at the beginning of the sweep in those scenarios. This allowed us to characterize the effects of historical recombination previous to the outset of the selective sweep. For each simulated population, the haplotypes, defined as the whole linkage group under selection, were classified in two groups: one group for haplotypes carrying the advantageous allele and one group for haplotypes with the neutral allele. Genetic diversity (measured as the average expected heterozygosity per bp) within the haplotypes carrying the advantageous allele and differentiation between the two groups of haplotypes (measured as the *F*_ST_) were calculated in 3 000 bp windows at increasing distance from the locus under selection.

### Real data application

*Medicago truncatula* is an annual, predominantly selfing species (Siol *et al.* 2008) of the legume family (Fabaceae), found around the Mediterranean Basin. We conducted a temporal survey in a population located in Cape Corsica (42° 58.406’ North, 9° 22.015’ East, 362 m.a.s.l.). Samples of around 100 pods were collected in 1987 and 2009 along three transects running across the population, with at least one meter distance between each pod collected in order to avoid over-sampling the progeny of a single individual. The seeds were stored in a cold room between collection year and 2011, when plants were replicated from seeds in standardized greenhouse conditions. Using these samples, we collected leaf material from 64 plants from 1987 and 96 plants from 2009 grown in greenhouse for DNA extraction. Between 100 mg and 200 mg of young leaves were ground in liquid nitrogen in a Qiagen Retsch Tissue Lyser (Qiagen N. V., Hilden, Germany) during 2×1 min at 30 Hz. The fine powder was mixed with 600 μl of pre-heated lysis buffer consisting of 100 mM Tris-HCl pH 8.0, 20 mM EDTA pH 8.0, 1.4 M NaCl, 2 %(w/v) CTAB, 1 %(w/v) PVP40 and 1 %(w/v) sodium bisulphite plus 8 μl of 10 mg/ml RNAse per sample added extemporaneously. After incubation for 20 min at 65 °C under medium shaking, 600 μl of chloroform were added and mixed with a vortex mixer. Each sample was centrifuged for 15 min at 10 000 g and 10 °C and the upper phase was transferred into a new tube and then mixed with 60 μl of 3 M sodium acetate and 600 μl of cold isopropanol. DNA was precipitated by another centrifugation (30 min, 15 000 g, 4 °C), rinsed with 300 μl of 70 % cold ethanol, centrifuged again (10 min, 15 000 g, 4 °C), dried for 10 min at room temperature and resuspended in 100 μl of sterile deionized water. The genotyping was performed using two SNP chips specifically developed for *M. truncatula* (Loridon *et al.* 2013). Out of the 1920 SNPs, 137 were located in genes encoding flowering time, 721 in other candidate genes, in particular genes involved in symbiosis, and 1062 at random positions of the genome. The 1920 SNPs were widespread across all eight linkage groups (Table A2).

The genotyping assays were achieved at GenoToul (Genomic Platform in Toulouse, France) and at the BioMedical Genomics Center (Minneapolis, University of Minnesota, USA) using GoldenGate Assay (Fan *et al.* 2006, 2003) and respectively Illumina’s VeraCode technology (Lin *et al.* 2009) or Bead Array technology. Data generated from the BeadXpressTM reader (384 SNP × 480 DNA) or BeadArray Reader (1536 SNP × 192 DNA) were analyzed as detailed in Loridon *et al.* (2013).

These data were analysed using the temporal genome scan described in this paper, providing estimates of the effective population size (from *F*_ST_) and the selfing rate (from *F*_IS_) of the population. Confidence intervals (CI) for those estimates were obtained by an approximate bootstrap procedure over loci (DiCiccio and Efron 1992). For loci with a global MAF higher than 0.05, we obtained a *p*-value for the test of homogeneity. In order to control for multiple testing, FDR estimates were quantified with *q*-values for each locus, following Storey (2002).

## Results

### Accuracy of *N*_e_ estimates

In neutral scenarios, estimates of effective population size derived from *F*_ST_ performed reasonably well, decreasing in the presence of partial selfing following the *N*_e_ theoretical expectation (Fig. 1a). Selfing also caused an increase of error in the estimation of *N*_e_. In the presence of selection, estimates of effective population size decreased with the strength of the selective coefficient of the causal mutation (Fig. 1b). As expected, effective population size estimates from *F*_C_ showed a clear bias in the presence of partial self-fertilization while the bias was negligible in estimates from *F*_ST_ (Fig. A3a).

**Figure 1.**
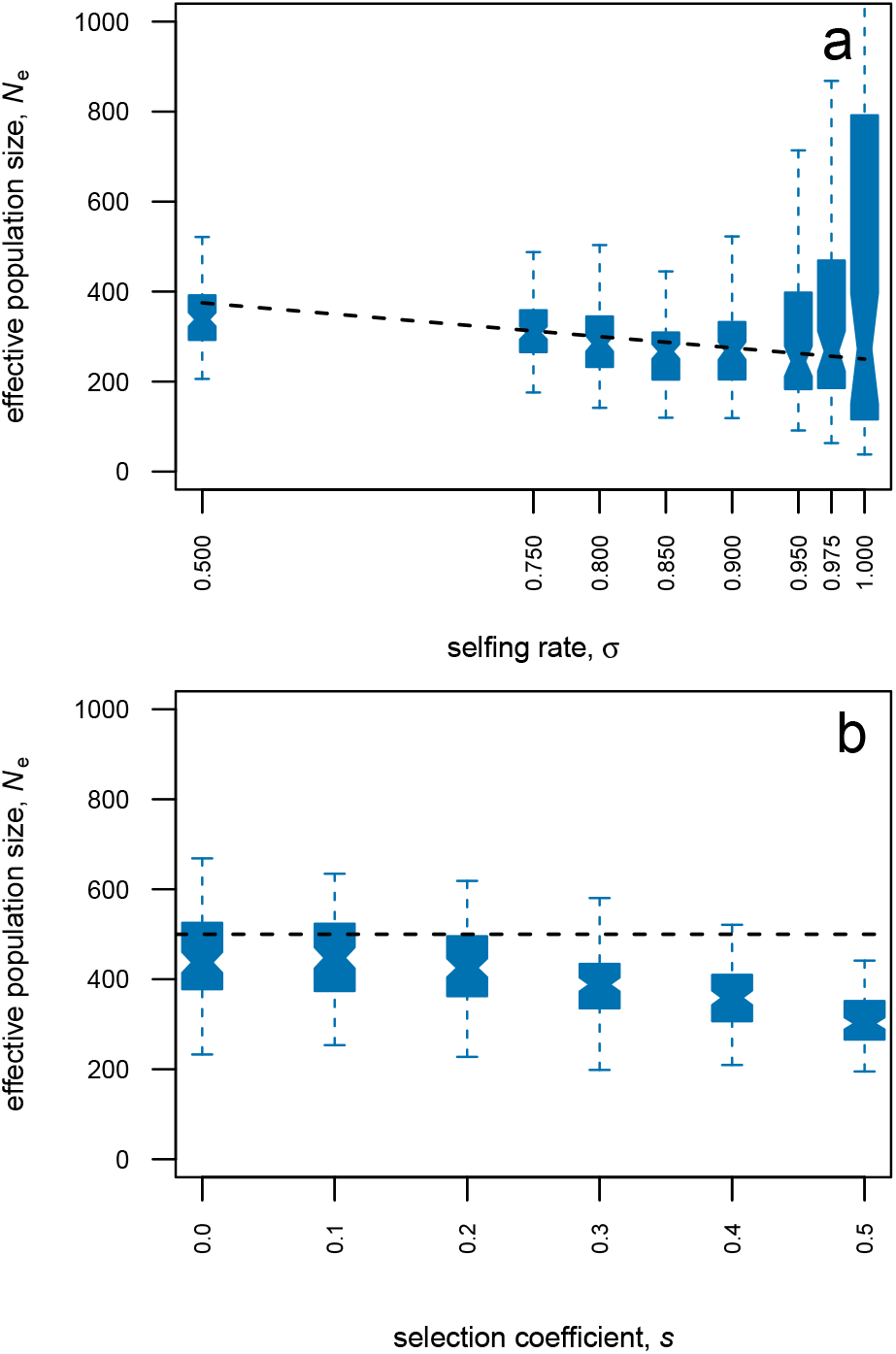
Effective population size estimates from temporal differentiation. Estimates of *N*_e_ from *F*_ST_ were obtained from simulated data of populations of *N* = 500 diploid individuals, sampled twice with *τ* = 25 generations between samples and selfing rate *σ*. **(a)** Neutral selfing population. **(b)** Outcrossing population under selection: a new advantageous mutation appears at generation *t* = 0 with coefficient of selection *s*. Samples are made of 50 diploid individuals genotyped at 10 000 biallelic markers (including the locus under selection), but only loci with a global MAF over 0.05 are used in the estimate (see details in Materials and Methods). Box-plot for estimates from 100 simulation replicates. Dashed line marks true effective population size, *N*_e_, in panel **(a)** and census size, *N*, in panel **(b)**.

### Footprint of selection

The temporal genome scan discriminated well between selected and neutral (on a separate linkage group) sites in most scenarios (Fig. 2). Yet, in the case of selection on new mutations, extreme selfing rate decreased the performance of the test (Fig. 2a). In scenarios of selection on standing variation the performance of the test was lower (Fig. 2b). In addition, the effect of selfing was more complex in the case of selection on standing variation. Moderate levels of selfing seemed to improve the discrimination capacity while the performance for high levels of selfing was similar to the one for outcrossing simulations. Only complete selfing reduced dramatically the discrimination capacity of the method.

**Figure 2.**
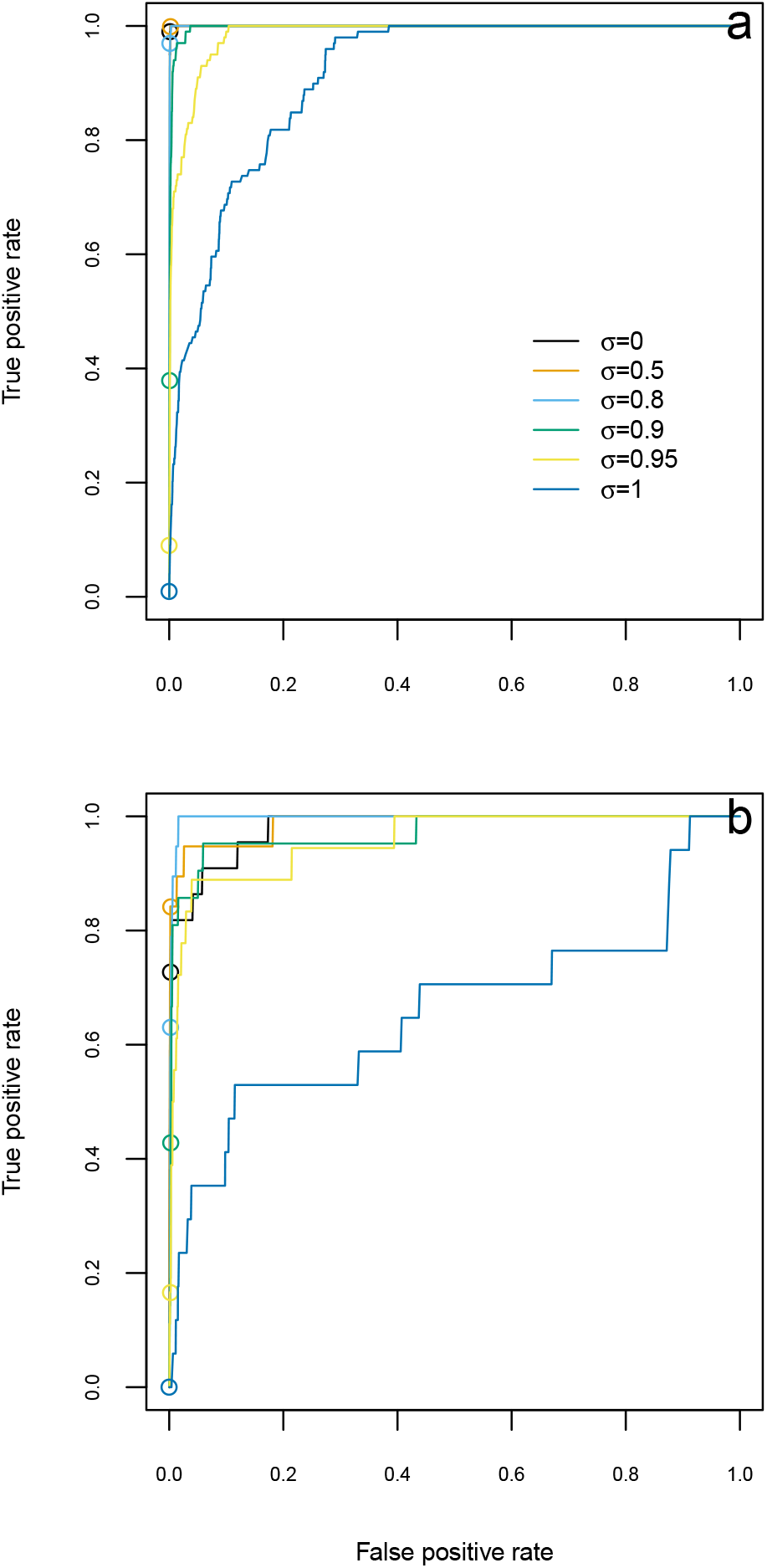
Power and false positive rate of the genome scan for increasing decision thresholds. Receiver operating characteristic curve is estimated from 100 replicates of simulated data of one population of *N* = 500 diploid individuals, sampled twice with *τ* = 25 generations between samples, selection coefficient *s* = 0.5 and selfing rate *σ*. **(a)** Selection on new mutation. **(b)** Selection on standing variation. Circles mark the values for positive threshold of *p*-value>0.001, which is the threshold used in Fig. 3.

These results, however, only consider the causal polymorphisms and completely independent neutral variants, ignoring linked sites that could have been subject to hitch-hiking. In practice, the signal for the detection of regions under selection comes mainly from those linked sites, which can create a local excess of outlier loci testing positive around the site under selection (even if the later is not in the data set). We therefore examined the footprint of selection at increasing distance from the advantageous allele, within a linkage group. As expected, the highest probability of positive test was at the locus under selection; then probability decreased with distance and reached very low values, similar to those for neutral loci on a separate linkage group (Fig. 3).

**Figure 3.**
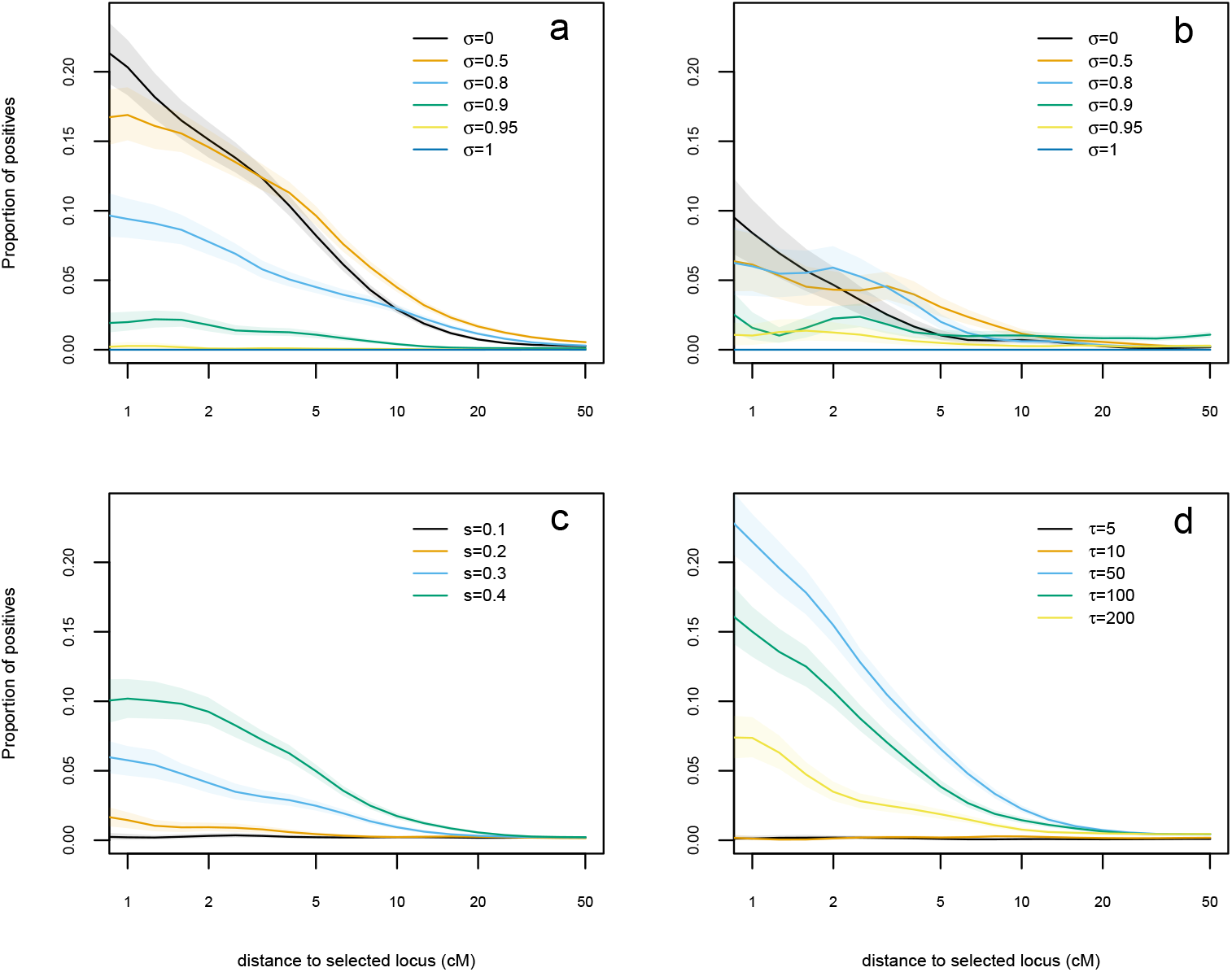
Selection footprint on the selected chromosome under different scenarios. Proportion of positive tests in function of distance to the locus under selection. Estimates obtained from 100 replicates of simulated data of one population of *N* = 500 diploid individuals, sampled twice with *τ* generations between samples and selection coefficient *s* and selfing rate *σ*. **(a)** Adaptation from new mutation, *τ* = 25, *s* = 0.5. **(b)** Adaptation from standing variation, *τ* = 25, *s* = 0.5. **(c)** Adaptation from new mutation, *τ* = 25, *σ* = 0. **(d)** Adaptation from new mutation, *s* = 0.5, *σ* = 0. Distance measured in centimorgans (cM). Coloured areas mark the 95% bootstrap interval.

The distance at which the hitch-hiking signal disappears depends on the scenario, being smaller for selection on scenarios of standing variation and larger for scenarios on new mutation, congruent with results by Hartfield and Bataillon (2020). Selfing rate increases distance at which there is a hitch-hiking effect (Fig. A5; Hartfield and Bataillon 2020). However, this did not translate into a signal from *F*_ST_ at larger distances, but to a reduction in power.

Under a scenario of adaptation from new mutations, the overall strength of the signal decreased with increasing selfing rate (Fig. 3a) and decreasing selection coefficient (Fig. 3c). The time sample interval also influenced the strength of the signal, with intermediate values being more favourable for the detection of outlier loci (Fig. 3d).

The results for the signal of hitch-hiking mirror those of the power to detect the causal site. In the case of selection on standing variation, there was an overall reduction of the signal compared to scenarios of selection on new mutation (Fig. 3a,b) except for highly selfing populations (*σ* ≥ 0.95). Nevertheless, as in the scenario of adaptation from new mutations, predominantly selfing populations had a weaker signal of selection than outcrossing populations. However, the strength of the signal for populations with intermediate levels of selfing (*σ* = 0.5, *σ* = 0.8) was similar or even higher than outcrossing populations (Fig. 3b).

The initial frequency of the advantageous allele and whether it is ancestral or derived also influenced the strength of the signal in predominantly selfing populations (*σ* = 0.95, Fig. 4). As expected, the lower the initial frequency, the stronger was the signal of selection, but this was modulated by the nature of the allele. With low initial frequencies, selection on the ancestral allele left a stronger signal. Symmetrically, the signal was stronger for derived alleles if the initial frequency was high. To understand these results, we examined the genetic diversity and the differentiation between the group of haplotypes carrying the advantageous mutation and the group not carrying it at the outset of the sweep. We found that this genetic differentiation decreases with the distance from the selected site, with higher differentiation for scenarios with stronger signal of selection (i.e. when the advantageous allele is ancestral with low initial frequency). Besides, ancestral alleles were associated to more diverse local haplotypes than derived alleles at the same frequency. These results show that no further recombination during the sweep is required to generate a local signal of selection.

**Figure 4.**
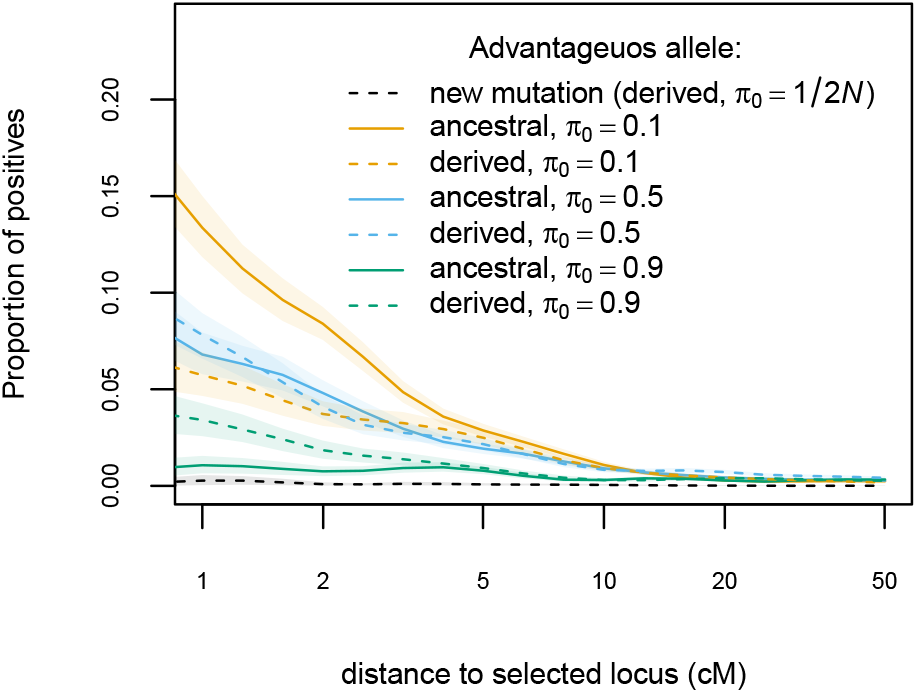
Influence of derived/ancestral state and initial frequency on selection footprint. Proportion of positive test in function of distance to the locus under selection. Estimates obtained from 100 replicates of simulated data of one population of *N* = 500 diploid individuals, sampled twice with *τ* = 25 generations between samples and selection coefficient *s* = 0.5. Adaptation on standing variation, selfing rate *σ* = 0.95.

### Medicago truncatula

From the 1920 SNPs genotyped, 1224 were polymorphic in the sample and 987 had a global MAF higher than 0.05. Genetic differentiation between time samples was large, 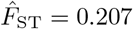 (95 % CI of 0.197 to 0.217) and so was the inbreeding coefficient, 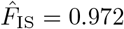 (95 % CI of 0.969 to 0.974). From those values, effective population size was estimated at 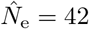 (95 % CI of 39 to 44) and selfing rate at 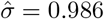 (95 % CI of 0.984 to 0.987). The test of homogeneity did not identify any SNP as a strong candidate for being under selection: the lowest *p*-value was 0.04 with a corresponding *q*-value of 0.48 (Fig. 5 and Table A2).

**Figure 5.**
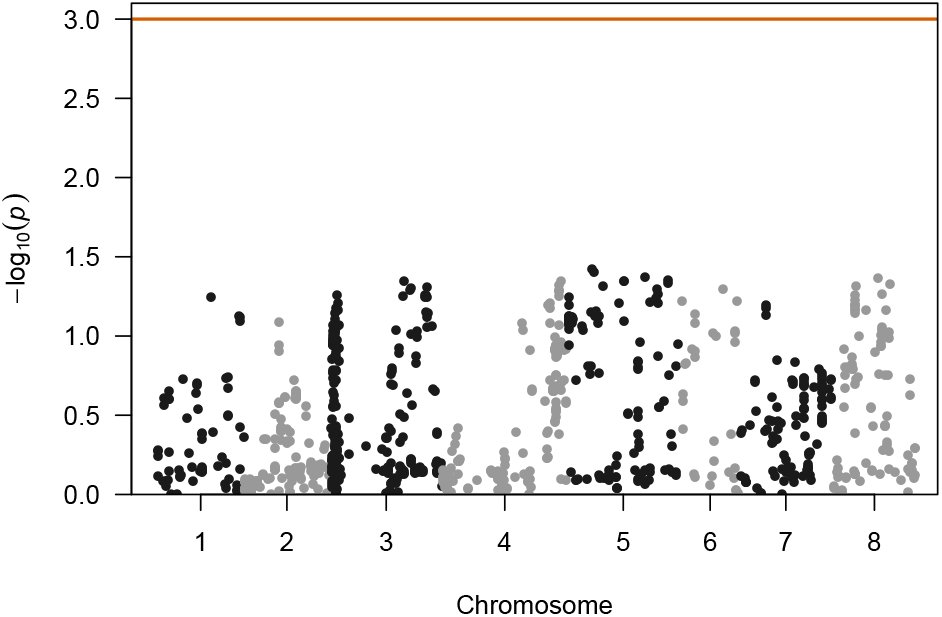
Genome-scan for selection based on temporal differentiation in cape Corsica *Medicago truncatula* population. − log_10_(*p*-value) of the simulation-based test of the null hypothesis that the locus-specific differentiation measured at each SNP is only due to genetic drift. Only SNP markers with MAF>0.05 and known position at the reference genome are shown. Red line marks *p*-value=0.001, the significance threshold used for figures 3 and 4.

## Discussion

### Estimating effective population size under selfing and selection

It is well known that self-fertilization reduces the effective size of populations. Our results show that Weir and Cockerham’s (1984) *F*_ST_ allows to measure the amount of drift between two temporal samples, even in the presence of selfing. However, the precision of the estimates diminishes with the rate of selfing. We hypothesize that it is the consequence of the effect of reduced effective recombination in selfing, which reduces the number of loci with independent evolutionary histories. This effect is most extreme for completely selfing populations, where the whole genome behaves as a single locus, reducing dramatically the information available in the data. In order to test this hypothesis, we estimated *N*_e_ for a similar set of simulations under an unrealistic model where each locus was simulated independently (appendix A2). On those simulations, the precision of *N*_e_ estimates was no longer reduced by selfing (Fig. A3b).

An alternative and common way to estimate *N*_e_ from temporal data uses the standardized variance in allele frequencies (*F*_C_, Nei and Tajima 1981). This was the approach originally proposed for the temporal genome scan by Goldringer and Bataillon (2004). However, estimates from *F*_C_ suffer from a bias (Fig. A3) because this approach assumes that the 2*n* gene copies (in a sample of *n* diploid individuals) are independent draws from the population gene pool, and uses a binomial distribution to model it (Waples 1989). However, in the case of partially or predominantly selfing populations, the two gene copies within an individual are not independent samples from the population, which explains the bias of the estimate. Using Weir and Cockerham’s (1984) *F*_ST_ estimate corrects for this bias.

Strong selection also reduces the effective population size by reducing the number of breeding individuals or increasing the variance of reproductive success among individuals (Robertson 1961; Santiago and Caballero 1995). This is reflected in the estimation of *N*_e_, which decreases with the strength of the selective coefficient of the mutation under selection in our simulations (Fig. 1b). Note that, in our case, this effect is not only driven by the local increase of temporal genetic differentiation around the site under selection. The strong selection considered in our simulations increases *F*_ST_ genome-wide, even in regions unlinked to the site under selection (e.g. distance of about 50 centimorgan or larger; Fig. A5). The combined effect of selfing (i.e. increased linkage disequilibrium) and selection produced very strong drift compared to a neutrally evolving population with the same census size (Fig. A5).

The effective population size estimated in Cape Corsica *M. truncatula* was extremely low, in agreement with the results from microsatellites data on the same population (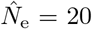, Jullien *et al.* 2019). However, such small *N*_e_ estimates might reflect other processes in addition to drift. For instance, gene flow into the studied population can increase temporal *F*_ST_ and bias *N*_e_ estimates (Jullien *et al.* 2019). The effects of the violation of model assumptions are discussed further below.

In our simulations we have considered that individual genotypes are available. However, in many temporal experiments, sequencing is performed for pools of individuals (Pool-seq; e.g. Franks *et al.* 2016) to reduce the costs. Estimating effective population sizes remains possible, using Pool-seq adapted *F*_ST_ and *F*_C_ estimates (Hivert *et al.* 2018; Jónás *et al.* 2016). However, since individual information is lost, these estimates cannot take into account the within individual allele identity correlation due to selfing. When studying a selfing species, we therefore recommend working with individuals genotypes. If Pool-seq is necessary due to budget limitations, we advise to follow a similar procedure as in Frachon *et al.* (2017) and reproduce seeds by selfing for one or more generations so that sequenced individuals are completely homozygous and can be treated as effectively haploid samples. However, this approach is likely to only give good results with predominantly selfing species (with a small proportion of heterozygous loci to be removed) and divides the amount of data by two.

### Limits imposed by selfing to temporal genome scans of selection

Our results show that the search for regions under selection with temporal *F*_ST_ genome scans in predominantley selfing populations can be challenging. In the case of complete selfing, the whole genome behaves as single locus and selection, if present, affects the whole genome. In partially selfing populations, the strength of the signal left by selection depends on whether the advantageous allele had the time to recombine with different genetic backgrounds. In a population with a selfing rate as high as *σ* = 0.95, outcrossing events are unlikely to occur during the sweep of a new beneficial mutation and produce no effective recombination if they happen between close relatives sharing the same homozygous genotype. As a results, like completely selfing populations, the selective sweep will reduce genetic diversity on the whole genome and will leave no local signal of selection.

On the contrary, when adaptation proceeds through standing variation, historical recombination can put an allele on different genetic backgrounds before it becomes advantageous and sweeps. In that case, the sweeping haplotypes are only similar among them and different from the haplotypes in the rest of the population around the site under selection. As the distance to the selected site increases, the diversity of the sweeping haplotypes raises and the differentiation with the other haplotypes segregating in the population decreases due to the action of historical recombination (Fig. A4). These differences along the genome create the local signal of selection, even if no further (effective) recombination occurs during the sweep.

Selection on standing variation is usually associated to a weak signal of selection (i.e. a soft sweep, Hermisson and Pennings 2005). Our results indeed confirm this expectation for outcrossing populations (compare Fig. 3a,b for *σ* = 0). Interestingly, for predominantly selfing populations, the situation is reversed. Yet, the strength of the local signal of selection, and therefore the power of the outlier test depends not only on the initial frequency of the advantageous allele, but also on its genetic background. Consider an advantageous derived allele at low frequency. Because of its low frequency, it is likely a young allele (Fig. A6) and few mutations will have accumulated around it. The selected haplotypes will therefore display low diversity and little differentiation from the deleterious haplotypes (Fig. A4). Thus, we expect that only few mutations will show a significant change in allele frequencies around the site under selection (e.g. Fig. A2a). On the other hand, if the ancestral allele (at low frequency) becomes advantageous, it will be likely to be on a very old lineage (since a high frequency derived allele is expected to be old). Being older, such allele has had more time for mutations to accumulate on both lineages, creating diversity that is unique to the haplotypes under selection. This will lead to more sites with a significant allele frequency change around the causal mutation (e.g. Fig. A2b). In the case of selection on an allele starting at high frequency, the situation is reversed. The lowest signal is for selection on high frequency ancestral alleles that not only will have a small change on allele frequency but also carry diversity common to the whole population. Finally, low frequency ancestral alleles might have the potential to create the strongest signal, but they are scarce in an equilibrium population (Fig. A6) so it is unlikely they represent a frequent case of sweep from standing variation.

Although temporal genome scans in predominantly selfing species are able to reveal the footprint of selection on standing variation, how prevalent is adaptation from pre-existing variation in selfing populations remains an open question. In the short-term, selection is most likely acting on standing variation (Barrett and Schluter 2008), but there are no specific predictions for selfing populations. Theoretical models predict an overall limitation of adaptation in selfers compared to outcrossers (Hartfield and Glémin 2016). However, selfers can hold high levels of cryptic genetic variation for polygenic traits, due to negative linkage disequilibrium. This diversity may allow a selfing population to adapt to changing conditions as quickly as outcrossing populations (Clo *et al.* 2020). Indeed, there is some evidence of selective sweeps in selfing species (Bonhomme *et al.* 2015; Huber *et al.* 2014). In agreement with this, the temporal genome scan in a predominantly selfing population of *A. thaliana* from a previous study showed footprints of selection (Frachon *et al.* 2017). Such positive results could have been favoured by the relatively large genetic variation of this population (Baron *et al.* 2015; Frachon *et al.* 2017). Based on the simulation results we present here, we can assume that this population has been adapting from standing variation over a short period of time (eight generations) rather than from many new mutations occurring during (or shortly before) the studied period.

The absence of evidence for selection in the *M. truncatula* dataset is manifest. It is reasonable to attribute this results to the extreme selfing rate estimate 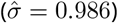 in this population, for which our simulation approach gives little hope to detect any signal of selection. Indeed, the same selfing lineages are observed throughout the study period of 22 generations (Jullien *et al.* 2019), which further suggest that almost no recombination occurs in the population. If selection, either on new or pre-existing mutations, has occurred, it has changed the frequencies of multilocus genotypes without leaving any local signal along the genome. This would have dramatically reduced the effective population size, which is consistent with our extremely low estimate 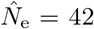. An examination of the distribution of *p*-values from the genome scan shows a departure from the expected distribution which could indicate an inappropriate null model (Fig. A7a). This deficit of low *p*-values means that the outlier test is too conservative because the null distribution of temporal *F*_ST_ is wider than the distribution observed genomewide. However, the QQ plots for simulated populations show that, as the selfing rate increases, the test progressively shifts from delivering an excess of low *p*-values to delivering a deficit of low *p*-values (Fig. A7b), as observed for *M. truncatula*. The present analysis cannot be conclusive about the presence of selection and the observed diversity changes can be the consequence of demographic processes, such as genetic exchanges with neighbouring populations, as discussed by Jullien *et al.* (2019).

### Model assumptions

The method and simulations presented here are based on simple models with one unstructured population, constant parameters through time and a single selective event. These models are useful to highlight the effects of inbreeding and to point out to some solutions for the problems posed by the presence of inbreeding. Real populations are complex systems and other demographic and selective processes are in action and they have consequences for the dynamics of the selective sweeps and the performance of the statistical methods.

The assumption that temporal data collected from the same geographical location belong to a single continuous population is probably wrong in many cases, even more so for predominantly selfing populations, where the spatial structure is usually very strong (Bonnin *et al.* 2001). Estimates of effective population size from temporal samples have been shown to be affected by population structure, often leading to underestimation (Gilbert and Whitlock 2015; Ryman *et al.* 2014). Such bias can make the tests conservative, aggravating the problem of detecting loci under selection. One way to address this issue is to perform demographic inference under more complex scenarios to be able to discriminate isolated from admixed populations, and jointly estimate drift, migration and selfing parameters (e.g. using approximate Bayesian computation as in Jullien 2019). Note, however, that some scenarios might enhance the signal of selection. Detection of sweeps from standing variation could be easier in populations with a larger historical effective size than in populations that had always been at a smaller constant size, because they benefit from higher historical recombination.

The presence of multiple loci under selection is also an important factor to consider, because strong selective interference is expected under selfing (Hartfield *et al.* 2017). Of particular interest is background selection which reduces the effective population size, an effect exacerbated by selfing (Nordborg 1997; Roze 2016). In addition to the reduction in diversity, purifying selection could also mimic the temporal signal of a selective sweep. Neutral alleles that are linked to less deleterious backgrounds can quickly rise to high frequencies (Cvijović *et al.* 2018). Recently, Johri *et al.* (2020) proposed using approximate Bayesian computation for joint inference of demography and purifying selection, which could lead to more appropriate null models for the detection of loci involved in adaptation. Further research is needed in this direction, it is unclear at this point the implications for temporal genome scans and partially selfing populations.

Future developments should not focus only on improving the inference of the null model. The classification of loci as neutral or into different selection categories can benefit from other information than just the genetic differentiation. For instance, reduction of genetic diversity around the locus under selection (e.g. Fig. A4) can add information about the presence and origin of the selection. Supervised machine learning methods have been shown to perform well and are promising tools for this type of task (reviewed by Schrider and Kern 2018). While these methods might improve the classification of loci under scenarios where there is some local footprint of selection, predominantly selfing population will remain a challenge as long as selective sweeps produce genome-wide hitch-hiking.

## Conclusions

Identifying regions under selection with a temporal genome scan can fail for several reasons. First, timing is paramount. A sample at the beginning and at the end of the selective sweep would be ideal for detecting selection. However, the start and duration of the sweep and the sampling times cannot be synchronized except for some experimental evolution studies. Frequency of the advantageous allele at the beginning of the sweep can also reduce the chances of capturing any signal. On top of that, selfing presents itself as a strong additional difficulty for this task, reducing the efficacy of selection (reduced *N*_e_) and extending hitchhiking effects that blur the distinction between neutral and selected regions. Nevertheless, scenarios of adaptation from standing variation can leave some signal of selection in highly selfing populations. Curiously, adaptation from new mutation under the same highly selfing scenarios leaves no signal, reversing the general expectation for the footprint of selection from “hard” and “soft” sweeps. Chance plays an important role in the success of a genome scan. Therefore, researchers should focus on the factors that can be controlled, such as the use of adapted methods to selfing species, to increase the probability of a positive outcome.

## Author contributions

Miguel Navascués, Laurène Gay, Joëlle Ronfort and Renaud Vitalis conceived and designed the research. MN and RV developed the temporal *F*_ST_ scan for partially selfing populations and the code implementing it. Arnaud Becheler and MN evaluated the performance of the method. Karine Loridon, LG and JR produced the *Medicago truncatula* data set. MN wrote the article with the help of all authors that critically reviewed and approved the text.

## Data and code accessibility

The method described in this work is implemented in DriftTest, a C program available at Zenodo (Navascués and Vitalis 2020). An R script (R Core Team 2018) to reproduce the simulations and analyses in this work is available at Zenodo (Becheler *et al.* 2018). SNP data for *Medicago truncatula* is available at Data INRAE (Gay 2020).

## Acknowledgements

This research was developed under the SelfAdapt project, funded by INRA metaprogramme “Adaptation of Agriculture and Forests to Climate Change” (ACCAF). A large part of the analyses presented in this work was performed on the CBGP HPC computational platform; we thank the platform manager, Alexandre Dehne Garcia, for being ever-helpful. We are grateful to Matteo Fumagalli, Christian Huber and two anonymous reviewers for their comments on this work. Version 4 of this preprint has been peer-reviewed and recommended by Peer Community In Evolutionary Biology (doi:10.24072/pci.evolbiol.100110)

## Appendices

## A1 Predominantly selfing species in project Baseline

Project Baseline will track the evolution of natural populations from more than 60 plant species in the next 50 years through seed collection (www.baselineseedbank.org, Etterson *et al.* 2016). Many of the species included are capable of self-fertilization and at least six of them reproduce predominantly by selfing: *Bromus diandrus* (Kon and Blacklow 1990), *Bromus tectorum* (Novak *et al.* 1991), *Elymus canadensis* (Sanders and Hamrick 1980), *Impatiens pallida* (Schemske 1978), *Stipa pulchra* (Larson *et al.* 2001) and *Triodanis biflora* (Goodwillie and Stewart 2013).

## A2 Simulation of independent loci

Additional simulations were performed to show the effect in the estimation of *N*_e_ of nonindependent sample of gene copies within individuals (i.e. *F*_IS_) without the effect of linkage disequilibrium due to selfing. The simulation approach is very similar to the simple drift model used to build null distribution of the test described in the main text. The main difference is that initial allele frequency is set as a fixed parameter. Simulations are used to generate temporal data at 10 000 biallelic loci per pseudo-observed dataset. Initial allele frequencies (*π*_0_) are fixed to 0.5 for all loci. Allele frequency *π* in generation *t* were simulated with a binomial distribution as *π_t_ ∼* B(2*N*_e_, *π_t−_*_1_)*/*2*N*_e_, where *N*_e_ is the effective population size in number diploid individuals, for generations *t ∈* [1, *τ*]. Thus, each locus is simulated independently from each other, ignoring the linkage among them that should have occurred due to selfing. Genotype counts in samples, ^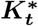^, at time *t* = 0 and *t* = *τ* are taken from a multinomial distribution, 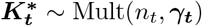, where *n_t_* is the sample size (in number of diploid individuals) at time *t* and *γ_t_*:

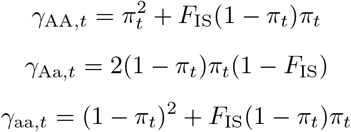

are the genotype frequencies in the populations in function of the inbreeding coefficient *F*_IS_, which is determined by the selfing rate *F*_IS_ = *σ/*(2 *− σ*). Simulations were performed with parameters values for *N*_e_ = 500 diploid individuals, *τ* = 25 generations, *n*_0_ = *n*_25_ = 50 diploid individuals and selfing rate, *σ*, had values of 0, 0.5, 0.75, 0.8, 0.85, 0.9, 0.925, 0.95, 0.975 or 1. Results are presented in Fig. A3.

**Table A1.** Notation.

| symbol | meaning |
| --- | --- |
| $\gamma_t$ | genotype frequencies in the population at time $t$ |
| $\pi_t$ | allele frequency (in the population) at generation $t$ |
| $\sigma$ | rate of reproduction by selfing in the population |
| $\tau$ | time in generations between the two samples |
| AA, Aa, aa | the three possible genotypes of a bi-allelic locus |
| $F_{ST}^l$ | observed differentiation statistic $F_{ST}$ at focal locus $l$ |
| $K_t$ | observed genotype counts in the sample at time $t$ |
| $K_t^*$ | simulated genotype counts in the sample at time $t$ |
| $k_t$ | observed count of the reference allele in sample at time $t$ |
| $N$ | census population size in number of diploid individuals |
| $N_e$ | effective population size in number of diploid individuals |
| $n_t$ | sample size (number of diploid individuals) at time $t$ |
| $p_t$ | frequency of the reference allele in the sample at generation $t$; $p_t = k_t/2n_t$ |
| $p_t^*$ | simulated frequency of the reference allele in the sample at generation $t$; $p_t = k_t/2n_t$ |
| $s$ | selection coefficient of the adaptive locus |
| $t$ | time, measured in generations, with $t = 0$ for first sample and $t = \tau$ for last sample |

**Table A2.** Genome scan results for *M. truncatula*. This table is provided in a supplementary file.

**Figure A1.**
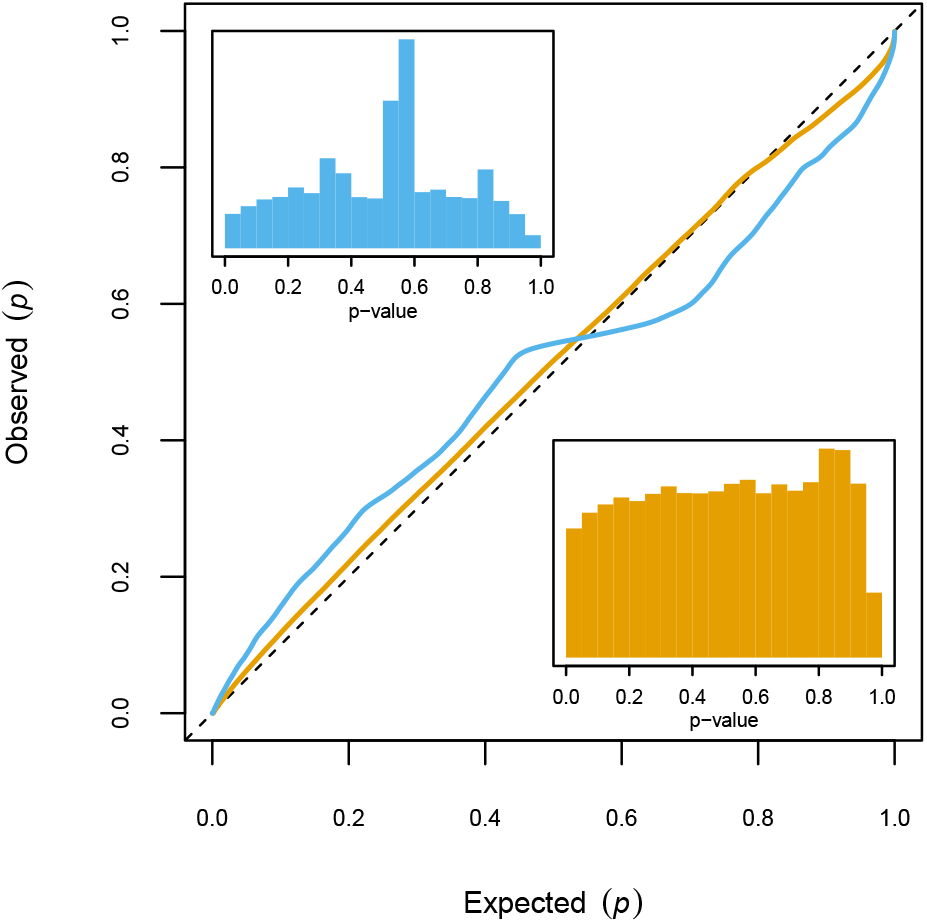
Effect of a filter on minor allele frequency (MAF) on the distribution of *p*-values. *p*-values calculated from 100 simulation replicates of a neutral outcrossing population of *N* = 500 diploid individuals, sampled twice with *τ* = 25 generations between samples. The blue histogram shows the distribution of *p*-value using all loci, with corresponding blue line in the QQ-plot. The orange histogram shows the distribution of *p*-value using loci with a minimum global allele frequency of 0.05, with corresponding orange line in the QQ-plot.

**Figure A2.**
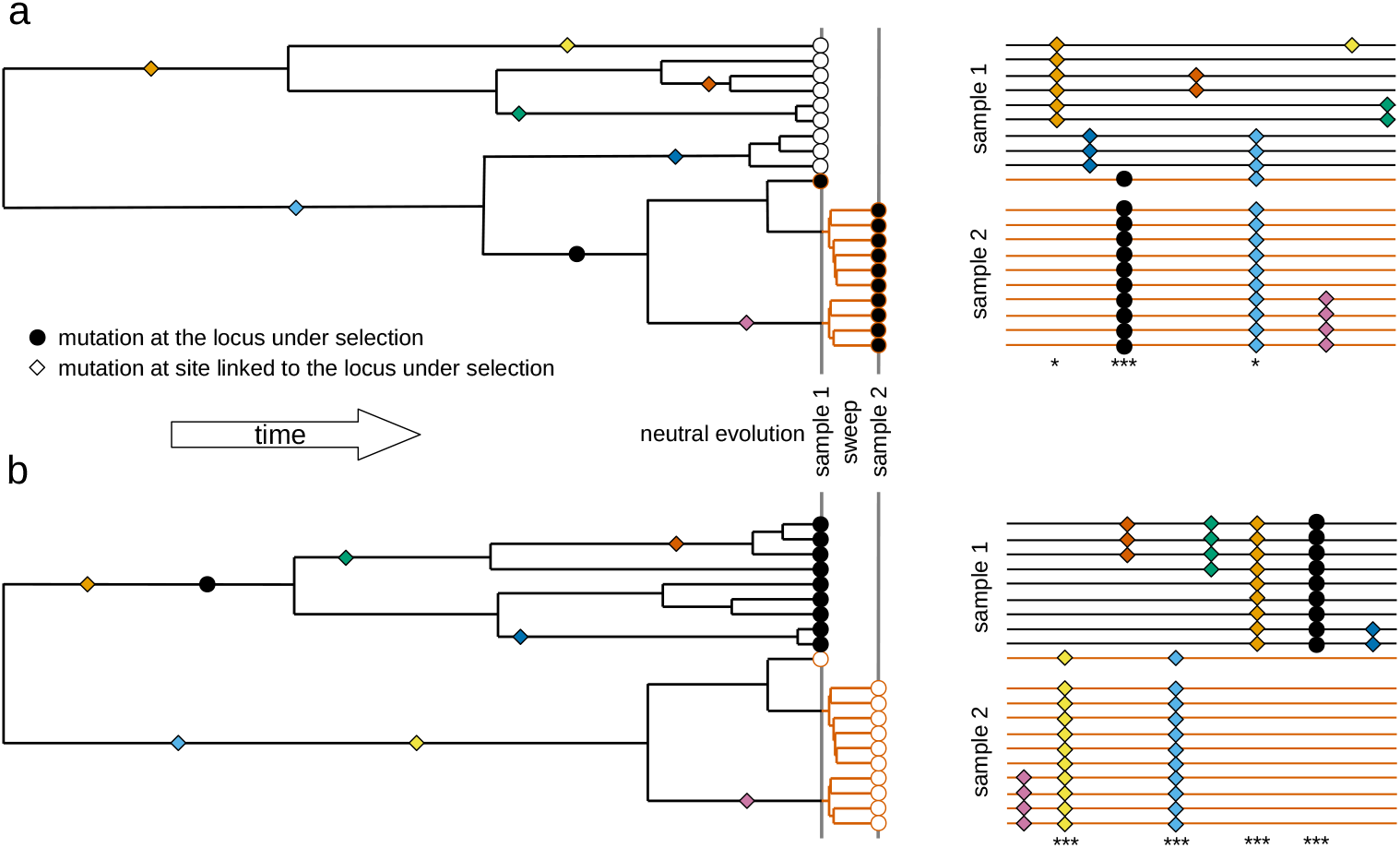
Schematic representation of the consequences of selection on standing variation in the temporal pattern of genetic diversity. Gene genealogies with samples carrying the derived allele represented with a circle filled in black and samples carrying the ancestral allele represented with a circle filled in white. Mutations are represented as coloured squares over the genealogy at their time of mutation; the site under selection is represented with a circle filled in black. Lineages of the genealogy subject to positive selection are represented in red. Next to each genealogy a schematic representation of the sequence alignment of the samples is presented with each sequence represented as a line at the same height that the corresponding sample of the genealogy. Polymorphisms are represented with the corresponding coloured squares in the genealogy, indicating the sequences that carry the derived alleles. Sequences carrying the allele under positive selection are represented with a red line. Asterisks signal polymorphic sites showing large (*) and extreme (***) changes in allele frequency between the two temporal samples. This figure represents only a small region around the locus under selection that did not recombine through the period considered (hence can be represented as a single genealogy). **(a)** Gene genealogy of two temporal samples taken just before a low frequency derived allele becomes advantageous and after the sweep. The mutation is young, as expected for a low frequency derived allele (Fig. A6), so the lineages carrying the allele that will become advantageous had little time to accumulate mutations and become distinctive from other haplotypes in the population. Most of the alleles on the haplotypes that swept and are found in the second sample on high frequency were already at high frequency in the first sample. **(b)** Gene genealogy of two temporal samples taken just before a low frequency ancestral allele becomes advantageous and after the sweep. The mutation is old, as expected for a high frequency derived allele (Fig. A6); thus, the split between the lineages carrying the ancestral and derived allele is very old. Several mutations have accumulated in both lineages, making them very distinctive. Several alleles on the haplotypes that swept and are found in the second sample on high frequency were at low frequency in the first sample, creating a strong local signal of selection around the selected site.

**Figure A3.**
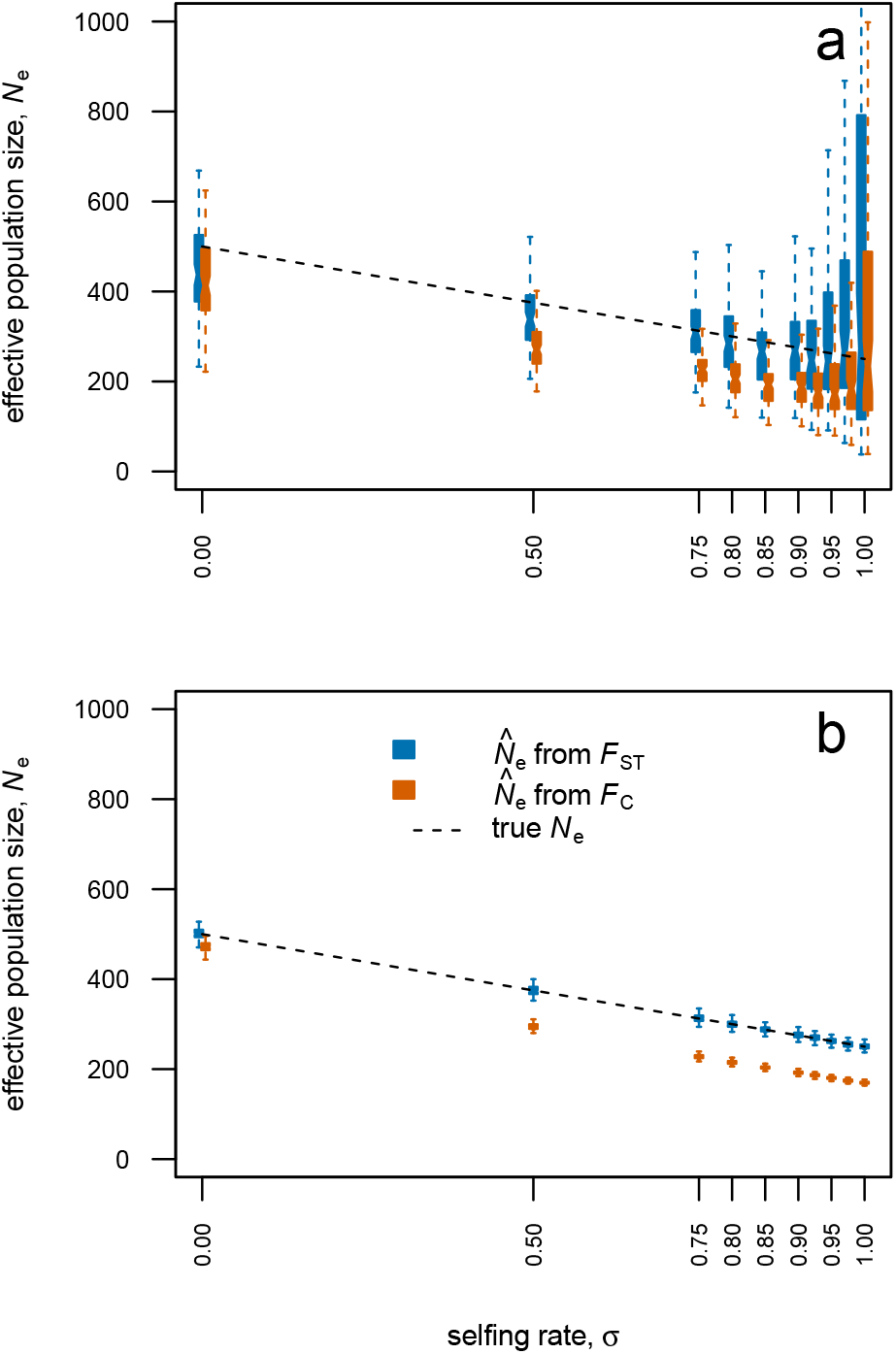
Effective population size estimates from temporal differentiation from *F*_C_ and *F*_ST_. Estimates obtained from simulated data of populations of *N* = 500 diploid individuals, sampled twice with *τ* = 25 generations between samples and 100 estimates at independent simulation replicates. Dashed line marks true 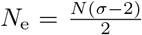. **(a)** Simulations performed with SLiM (Messer 2013) as described in the main text and presented partially in Fig. 1. **(b)** Simulations performed with a simple model of independent loci with initial allele frequency of 0.5, as described in appendix A2.

**Figure A4.**
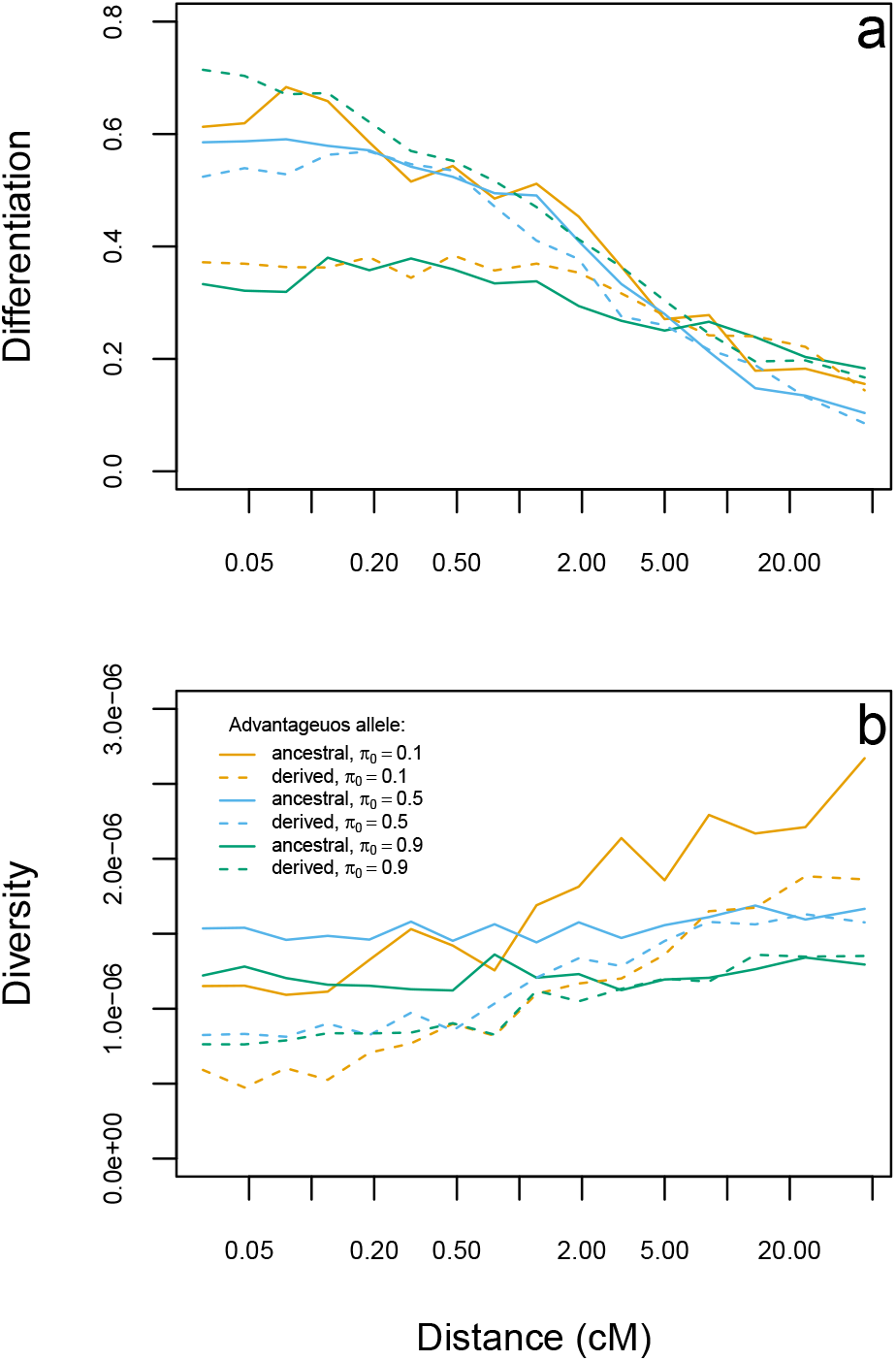
Effect of recombination occurring before the selective sweep on differentiation and diversity of the haplotypes carrying the advantageous allele. Genetic patters found in a simulated population of *N* = 500 diploid individuals reproducing predominantly by selfing (*σ* = 0.95) after 20*N*_e_ generations. **(a)** Differentiation is measured as the *F*_ST_ between haplotypes carrying the advantageous allele and those carrying the alternative allele. **(b)** Diversity is measured as the average heterozygosity per bp. Both differentiation and diversity were calculated on 3 000 bp windows at different distances to the loci under selection. The mean from 100 simulation replicates is shown.

**Figure A5.**
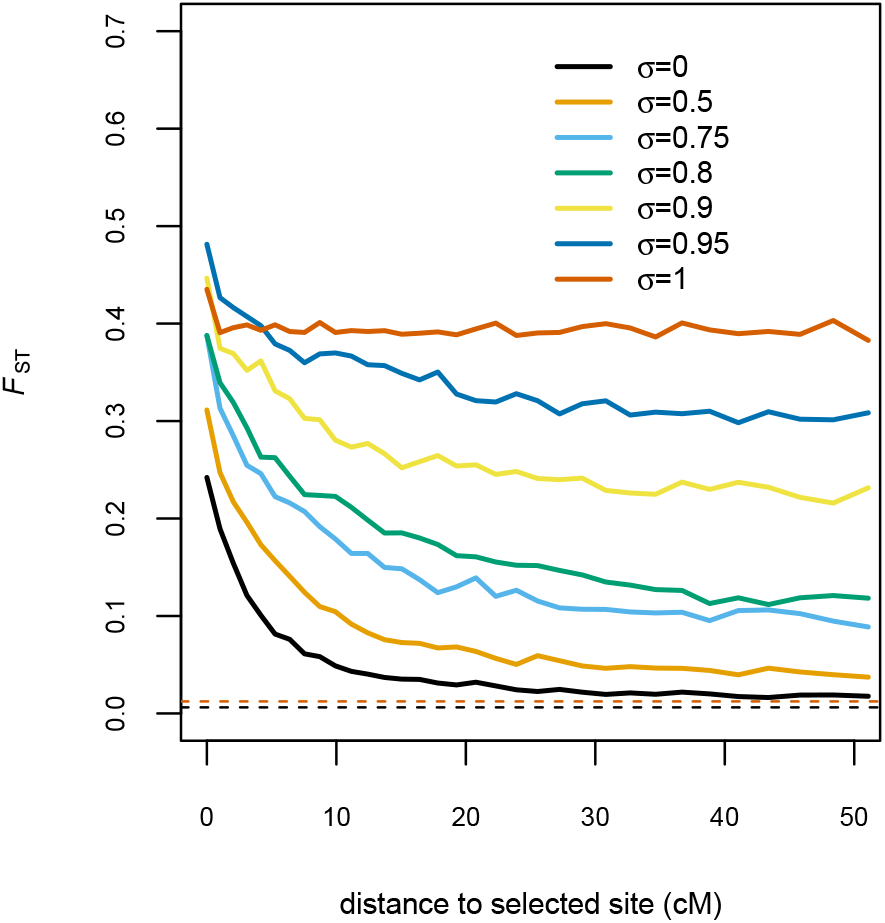
Genetic differentiation as a function of the distance to locus under slection. Estimates of *F*_ST_ (mean across 100 simulation replicates) from polymorphisms in a window of size 1 000Kb (*≈*1cM) at different distances to locus under selection. Simulation of populations of *N* = 500 diploid individuals, sampled twice with *τ* = 25 generations between samples and different rates of selfing, *σ*. A new advantageous mutation appears at generation *t* = 0 with coefficient of selection *s* = 0.5. Samples are made of 50 diploid individuals genotyped at 10 000 biallelic markers (including the locus under selection). For reference, expected *F*_ST_ values for *σ* = 0 (black) and *σ* = 1 (red) are shown with dashed lines.

**Figure A6.**
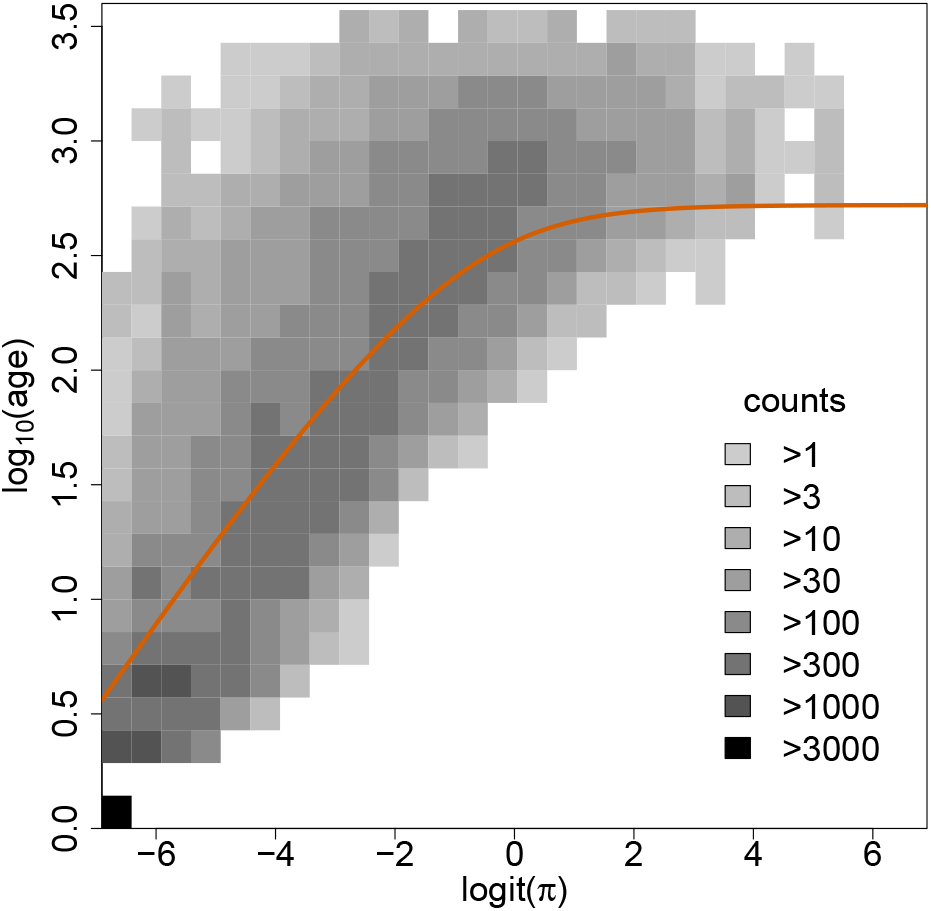
Relationship between frequency (*π*) and age of mutation. Bidimensional histogram for the frequency and age (in generations) of all mutations in a simulated population of *N* = 500 diploid individuals reproducing predominantly by selfing (*σ* = 0.95) after 20*N*_e_ generations. Theoretical expectation of age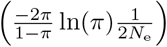 is shown with an orange line(Kimura and Ohta 1973).

**Figure A7.**
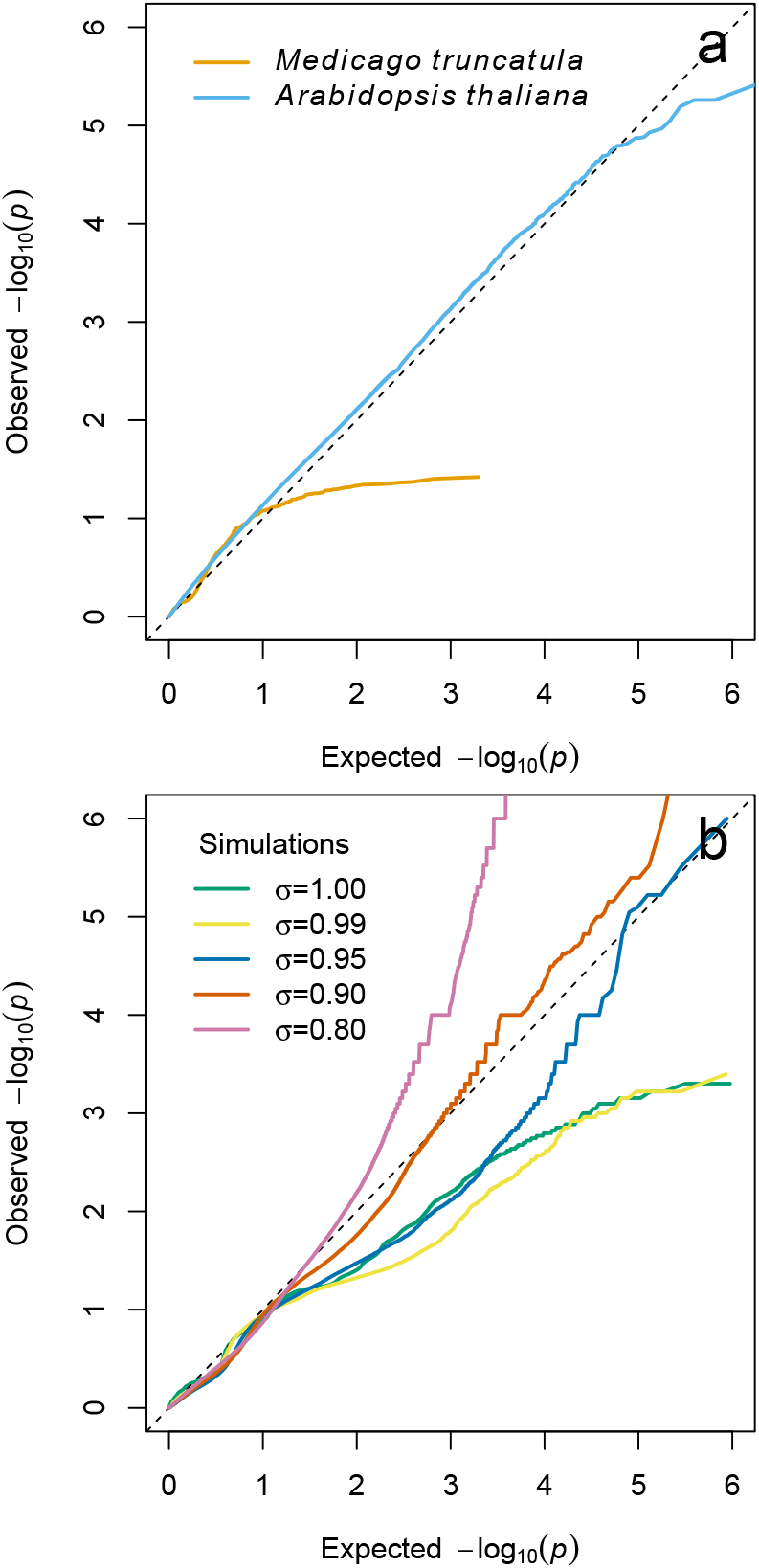
Departure from expectations of the *p*-value distribution for *Medicago truncatula* genome scan. **(a)** QQ plots for *Medicago truncatula* (this study) and *Arabidopsis thaliana* (Fig. S21 Frachon *et al.* 2017). For *M. truncatula*, *p*-values estimated for 987 SNP loci that were polymorphic in the sample with a global minor allele frequency higher than 0.05 (Table A2). Null model considered *N*_e_ = 42 and *σ* = 0.99. **(b)** QQ plots for simulations under a scenario of adaptation from new mutations in a population of *N* = 500 diploid individuals sampled twice with *τ* = 25 generations between samples and different rates of selfing, *σ*. A new advantageous mutation appears at generation *t* = 0 with coefficient of selection *s* = 0.5. Samples are made of 50 diploid individuals genotyped at 10 000 biallelic markers (including the locus under selection). The plot consider the distribution of *p*-values from 100 simulation replicates.

